# Follow-up investigation and detailed mutational characterization of the SARS-CoV-2 Omicron variant lineages (BA.1, BA.2, BA.3 and BA.1.1)

**DOI:** 10.1101/2022.02.25.481941

**Authors:** Qussai Abbas, Alexey Kusakin, Kinda Sharrouf, Susan Jyakhwo, Aleksey S Komissarov

## Abstract

Aided by extensive protein mutations, the SARS-CoV-2 Omicron (B.1.1.529) variant overtook the previously dominant Delta variant and rapidly spread around the world. It was shown to exhibit significant resistance to current vaccines and evasion from neutralizing antibodies. It is therefore critical to investigate the Omicron mutations’ trajectories. In this study, a literature search of published articles and SARS-CoV-2 databases was conducted, We explored the full list of mutations in Omicron BA.1, BA.1.1, BA.2, and BA.3 lineages. We described in detail the prevalence and occurrence of the mutations across variants, and how Omicron differs from them. We used GISAID as our primary data source, which provides open-access to genomics data of the SARS-CoV-2 virus, in addition to epidemiological and geographical data. We examined how these mutations interact with each other, their co-occurrence and clustering. Our study offers for the first time a comprehensive description of all mutations with a focus on non-spike mutations and demonstrated that mutations in regions other than the Spike (S) genes are worth investigating further. Our research established that the Omicron variant has retained some mutations reported in other SARS-CoV-2 variants, yet many of its mutations are extremely rare in other variants and unique to Omicron. Some of these mutations have been linked to the transmissibility and immune escape of the virus, and indicate a significant shift in SARS-CoV-2 evolution. The most likely theories for the evolution of the Omicron variant were also discussed.

## Introduction

Severe acute respiratory syndrome coronavirus 2 (SARS-CoV-2) is the causative agent of COVID-19, which emerged in December 2019 and it continued to evolve, creating different variants of progressively increased transmissibility between humans [1]. SARS-CoV-2 is an enveloped, positive-sense, single-stranded RNA virus with a genome size of about 30,000 base pairs of genus Betacoronavirus [2]. Genetic lineages of SARS-CoV-2 have been emerging and circulating the world since the beginning of the COVID-19 pandemic. Globally, various COVID-19 variants exist and some may become a concern to public health. Consequently, WHO designated SARS-CoV-2 variants of concern (VOCs) based on their high transmission rate, potential immune evasion, unusual epidemiological properties, or adverse impact on diagnostics and therapeutics. WHO also designated SARS-CoV-2 variants of interest (VOIs), depending on the fact that these variants have specific genetic markers, which are predicted or known to affect virus characteristics. So far, there are five main VOCs, Alpha (B.1.1.7), Beta (B.1.351), Gamma (P.1), Delta (B.1.617.2), and Omicron (B.1.1.529) [3]. There are also three VOIs, Lambda (C.37), Mu (B.1.621), and AY.4.2 [4]. Further information about VOCs and VOIs are shown in Supplementary Table 2.

The omicron variant (B.1.1.529) was first detected in South Africa and was reported to WHO on the 24th of November 2021 and designated as VOC on the 26th of November 2021 [4]. As of the 7th of January 2022, this variant has been confirmed in 135 countries [5], and the infected cases are the highest in the United Kingdom followed by the USA [6].

By comparing with the reference SARS-CoV-2 Wuhan-Hu-1 genome, there are more than 50 mutations in the genome of Omicron variant compared to other variants of concern. Substitution and deletion mutations are common in SARS-CoV-2 variants, but the insertion mutation (ins214EPE) has been observed for the first time in the Omicron lineages [7].

Several of these mutations are shared with previous VOCs, and some of them are associated with the increased ability to bind to the ACE2 receptor protein or involved in host immune system invasion [8–11].

Omicron variant possesses comparable binding affinity to human ACE2 in comparison with the wild type SARS-CoV-2, but much weaker binding affinity than Delta variant [12]. Recent study suggests that new mutations in Omicron variant contribute to transmission advantages, immune escape, and novel spike functionality [13, 14]. As of early February 2022, 8,609,048 SARS-COV-2 virus sequences have been submitted to the GISAID database [15]. Out of them 1,622,178 sequences are for the Omicron variant. Analyzing that many sequences is a bit of a technical challenge. World map epidemiology and geographic trajectories of Omicron mutations are available on the Nextstrain website [16].

### Various nomenclatures for Omicron

Omicron corresponds to Pango lineage B.1.1.529 and includes descendent Pango lineages BA.1, BA.1.1, BA.2 and BA.3 [17]. The Nextstrain project has assigned Omicron the clade identifier “21M” and inside this group assigned “21K” for “BA.1” and “21L” for “BA.2” [18]. The GISAID project has assigned it the clade identifier “GRA” [15].

### Omicron lineages

The three lineages of the Omicron variant (BA.1, BA.2, and BA.3) were first detected at approximately the same time and from the same place.

- BA.1 lineage was first identified on the 11th of November 2021, (Africa / Botswana / South East / Greater Gaborone / Gaborone), Accession ID on GISAID is [EPI ISL 6640916].
- BA.2 lineage was first identified on the 17th of November 2021, (Africa / South Africa / Gauteng / Tshwane), Accession ID on GISAID is [EPI ISL 6795834.2].
- BA.3 lineage was first identified on the 18th of November 2021, (Africa / South Africa / North West), Accession ID on GISAID is [EPI ISL 7605713].
- BA.1.1 is a BA.1 sub-lineage, and it differs from BA.1 only in one mutation in spike protein (R346K), which also occurs in the Mu variant. The number of BA.1.1 cases started to increase since the 8th of December 2021.

BA.1 is highly resistant to antiviral drugs and vaccine-induced humoral immunity. As well as, the Spike protein of BA.1 is less efficiently cleaved by furin and less fusogenic than those of Delta and an ancestral SARS-CoV-2 belonging to the B.1.1 lineage [19, 20]. Additionally, the pathogenicity of BA.1 is attenuated when compared to Delta and an ancestral B.1.1 virus [19, 20]. When compared to BA.1, Yamasoba et al. found that the BA.2 lineage has a higher effective reproduction number, higher fusogenicity, and higher pathogenicity, which supports the statistical findings that BA.2 has a 1.4 times higher effective reproductive number than BA.1. The team of Yamasoba et al. also showed that BA.2 is resistant to the humoral immunity induced by BA.1 [20]. In addition, the Omicron lineages (BA.1.1, BA.2, and BA.3) are likely more transmissible than omicron (BA.1) and Delta [21].

### The origin of the Omicron variant have been a subject of controversy

All viruses evolve over time, mutate, and new variants may occur with an increase in transmissibility, or an increase in virulence. Some variants may diverge early from other strains. The ability of viruses to adapt to new hosts and environments is enormously dependent on their capacity to generate de novo diversity in a short period. Spontaneous mutation rates vary among viruses. It is well established that RNA viruses mutate faster than DNA viruses, single-stranded viruses mutate faster than double-stranded viruses, and genome size appears to correlate negatively with mutation rate [22]. Omicron is so different from the millions of SARS-CoV-2 genomes that have been shared publicly that pinpointing its closest relative is difficult. Thus, it is crucial to identify the origin of the Omicron virus variant and how this heavily mutated variant develop.

There are mainly three explanations for how the Omicron variant may appear (Fig 1). The first theory proposes that Omicron could be developed and significantly mutated in an immune-suppressed patient, such as someone with HIV(Fig 1A). Previous research has revealed that immunocompromised individuals can serve as reservoirs for novel viral variants, such as human norovirus and influenza [24, 25]. Because South Africa has the world’s largest HIV epidemic, some experts speculated that Omicron originated there because it was first discovered there. The idea is bolstered by the sequencing of SARS-CoV-2 samples from some chronically infected patients, and the evidence is supported by the fact that persistent SARS-CoV-2 infections in immunocompromised patients can trigger the accumulation of an unusually high variety of mutations with potential relevance at both biological and epidemiological levels [26–29]. A recent study by Kemp, S. et al. found that chronic infection with SARS-CoV-2 led to viral evolution and impaired sensitivity to neutralizing antibodies in an immunosuppressed individual treated with convalescent plasma [30]. The study of Bazykin G. et al. reported an accumulation of 18 mutations de novo in the coronavirus samples from an immunodeficient patient (a lymphoma patient) [29]. These mutations accumulated through a 132-day observation period, which is exceeding the average rate of evolution of SARS-CoV-2. Interestingly, S:ΔH69/ΔV70 and S:Y453F mutations (the ΔF combination) were responsible for the bulk of positive samples. Y453F affects the receptor-binding domain (RBD), possibly increasing human angiotensin-converting enzyme 2 (hACE2) binding [31]. It allows immune escape from monoclonal antibodies and polyclonal sera. ΔH69/ΔV70 is also involved in the evasion of neutralising antibodies [30]. In other similar studies conducted by Borges V. et al. [28] and Karim F. et al. [27] it has been observed a concurrent accumulation of mutations de novo in the SARS-CoV-2 genome of immunocompromised patients (non-Hodgkin lymphoma patient under immunosuppressive therapy and advanced HIV patient with antiretroviral treatment failure, respectively) with a persistent SARS-CoV-2 infection over at least 6 months. The emerged mutations are potentially associated with immune evasion and/or enhanced transmission, primarily targeting the SARS-CoV-2 key host-interacting protein and antigen. Recent research has also demonstrated that immunocompromised paediatric and young adult patients are susceptible to prolonged viral infections with prolonged infectious virus shedding and mutation accumulation. The reported results support a potential correlation between host immune response and the emergence of viral variants, which may have the potential to escape antibody neutralisation [26]. These findings provide support to the hypothesis of intra-host evolution as one mechanism for the emergence of SARS-CoV-2 variants with immune evasion properties.

**Fig 1.**
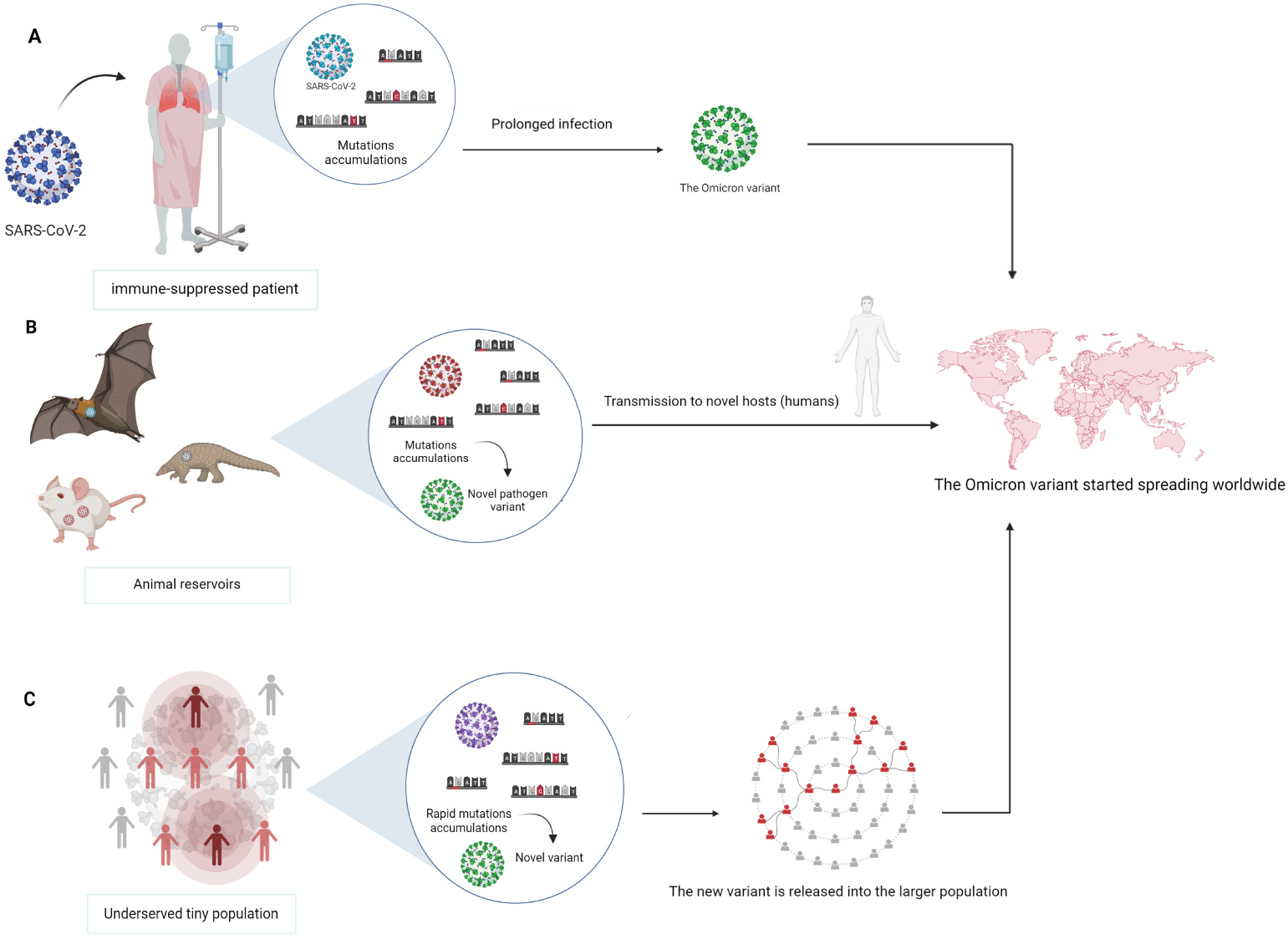
The three most likely scenarios for how the Omicron variant appeared. A: Omicron could have developed and significantly mutated in an immune-suppressed patient with prolonged viral infections. B: The Omicron variant might have evolved in non-human reservoirs, such as animal source, and as of late spread from it to humans. C: The virus might began circulating and changing in underserved places with a tiny population, where it had the opportunity to mutate rapidly, it could then have been released into the larger population, where it could have spread to different groups. Created with BioRender [23].

Alternatively, the Omicron variant might have evolved in non-human reservoirs, such as animal source, and as of late spread from it to humans(Fig 1B). Researchers have recently found evidence to support omicron’s potential origin in mice. The result of the team of professor Jianguo Xu suggests that omicron variant may evolve in mouse host. They discover 5 mouse-adapted mutation sites (K417, E484, Q493, Q498, and N501) in viral S protein suggesting mouse as intermediate host [32]. During evolution in mice, omicron adapted in host body by acquiring amino acid mutation in spike protein that enhanced binding with mouse ACE2 receptor [33, 34]. A recent study suggests that the Omicron variant has been around for much longer than predicted, given its close relationship to the Alpha variant [35].

Instead, the virus may have began circulating and changing in underserved places with a tiny population, where it had the opportunity to mutate rapidly in comparison to variations outside of that bubble (Fig 1C). It could then have been released into the larger population, where it could have spread to different groups. But that begs the issue of where Omicron’s predecessors were for all of that period, and is there someplace isolated enough for this sort of virus to transmit for that long without resurfacing in other places.

A recent study proposed that the Omicron “ins214EPE” insertion could have evolved in a co-infected individual. The study suggested that the nucleotide sequence encoding for ins214EPE could have been acquired by template switching involving the genomes of other viruses that infect the same host cells as SARS-CoV-2 or the human transcriptome of host cells infected with SARS-CoV-2 [7].

### Characterization of Omicron variant mutations

A vast and unprecedented number of mutations has accumulated in the Omicron lineages (BA.1, BA.2, and BA.3). These mutations spread over the genome, including Spike, and non-Spike regions. Some of these mutations are not unique to the Omicron variant; rather, some can be found in other VOCs or VOIs. Out of the 82 mutations, 24 mutations were found to be present in at least one additional variant. Consequently, 58 mutations have not been identified in any VOCs or VOIs, which make them “Omicron specific mutations”. In fact, the vast majority of mutations outside the Spike protein have not been identified in the previous VOCs or VOIs. After the investigation on the first occurrence place/time of each mutation (Table 1), it was revealed that several mutations appeared for the first time in North America (24 mutations). Others appeared for the first time in Asia, Europe, Africa, South America, or Oceania (19, 18, 12, 5, and 3 mutations, respectively). Interestingly, the bulk of these mutations first appeared in the first four months of 2020 (60 mutations in total).

**Table 1.** Omicron variant mutations prevalence and occurrence across variants. NTD: N-terminal domain; RBD: receptor-binding domain; RBM: receptor-binding motif; PBCS: poly-basic cleavage site; SD1/SD2: subdomain 1/subdomain 2; CTD: C-terminal domain.

| Mutation | Omicron lineage with the mutation | SARS-CoV-2 variants sharing the mutation with omicron variant (VOC, VOI) | First occurrence place | Collection date | Accession ID of the first detected viral sequence with the mutation (GISAID) |
| --- | --- | --- | --- | --- | --- |
| **NSP1** |  |  |  |  |  |
| S135R | BA.2 and BA.3 | — | Asia / Armenia | 2020-08-03 | EPI_ISL_1854612 |
| **NSP3** |  |  |  |  |  |
| T24I | BA.2 | — | Asia / Saudi Arabia / Makkah | 2020-04-06 | EPI_ISL_437704 |
| K38R | BA.1 and BA.1.1 | — | North America / Canada / Ontario | 2020-02 | EPI_ISL_418325 |
| G489S | BA.2 and BA.3 | — | Asia / China / Guangdong / Dongguan | 2020-01-26 | EPI_ISL_428466 |
| S1265del | BA.1 and BA.1.1 | — | South America / Brazil / Sao Paulo / Sao Jose dos Campos | 2020-03-09 | EPI_ISL_875545 |
| L1266I | BA.1 and BA.1.1 | — | South America / Brazil / Sao Paulo / Sao Jose dos Campos | 2020-03-09 | EPI_ISL_875545 |
| A1892T | BA.1 and BA.1.1 | — | North America / USA / New York / Erie County | 2020-04-17 | EPI_ISL_1009376 |
| **NSP4** |  |  |  |  |  |
| L264F | BA.2 | — | Europe / Italy / Lombardy | 2020-02-23 | EPI_ISL_542227 |
| T327I | BA.2 and BA.3 | — | Asia / China / Hubei / Wuhan | 2019-12-30 | EPI_ISL_402130 |
| L438F | BA.2 | — | North America / USA / Washington | 2020-03-13 | EPI_ISL_430895 |
| T492I | BA.1, BA.2, BA.1.1, and BA.3 | Delta, Mu, Lambda [36] | Asia / Georgia | 2020-02-28 | EPI_ISL_2896261 |
| **NSP5** |  |  |  |  |  |
| P132H | BA.1, BA.2, BA.1.1, and BA.3 | — | North America / USA / Texas / Houston | 2020-07-20 | EPI_ISL_5073555 |
| **NSP6** |  |  |  |  |  |
| A88V | BA.3 | — | Europe / Poland / Zachodniopomorskie | 2020-01-14 | EPI_ISL_2631277 |
| S106del | BA.1, BA.2, BA.1.1, and BA.3 | Alpha, Beta, Gamma [36] | North America / USA / New York | 2020-01-28 | EPI_ISL_3364539 |
| G107del | BA.1, BA.2, BA.1.1, and BA.3 | Alpha, Beta, Gamma [36] | North America / USA / New York | 2020-01-28 | EPI_ISL_3364539 |
| F108del | BA.1, BA.2, BA.1.1, and BA.3 | Alpha, Beta, Gamma [36] | North America / USA / New York | 2020-01-28 | EPI_ISL_3364539 |
| I189V | BA.1 and BA.1.1 | — | Europe / Ireland | 2020-03-10 | EPI_ISL_5248210 |
| **NSP12 (RdRp)** |  |  |  |  |  |
| P323L | BA.1, BA.2, BA.1.1, and BA.3 | Alpha, Beta, Gamma, Delta, Mu, Lambda [36] | Africa / Niger / Niamey | 2020-01-01 | EPI_ISL_3804266 |
| **NSP13** |  |  |  |  |  |
| R392C | BA.2 | — | Asia / China / Hubei / Wuhan | 2020-01-26 | EPI_ISL_493181 |
| **NSP14** |  |  |  |  |  |
| I42V | BA.1, BA.2, BA.1.1, and BA.3 | — | Asia / United Arab Emirates | 2020-03-15 | EPI_ISL_435127 |
| **NSP15** |  |  |  |  |  |
| T112I | BA.2 | — | North America / USA / Florida | 2020-03-19 | EPI_ISL_1379452 |
| **Spike protein (NTD)** |  |  |  |  |  |
| T19I | BA.2 | — | North America / USA / Colorado | 2020-03-17 | EPI_ISL_5439308 |
| L24S | BA.2 | — | North America / USA / Washington | 2020-03-17 | EPI_ISL_570806 |
| P25del | BA.2 | — | South America / Brazil / Rio de Janeiro | 2020-04-17 | EPI_ISL_1691459 |
| P26del | BA.2 | — | South America / Brazil / Rio de Janeiro | 2020-04-17 | EPI_ISL_1691459 |
| A27del | BA.2 | — | South America / Brazil / Rio de Janeiro | 2020-04-17 | EPI_ISL_1691459 |
| A67V | BA.1, BA.3, and BA.1.1 | Eta [37] | Europe / Spain / Madrid / Madrid | 2020-03-12 | EPI_ISL_779718 |
| H69del | BA.1, BA.3, and BA.1.1 | Alpha, Eta [37] | Asia / Thailand / Nonthaburi | 2020-01-05 | EPI_ISL_434692 |
| V70del | BA.1, BA.3, and BA.1.1 | Alpha, Eta [37] | Asia / Thailand / Nonthaburi | 2020-01-05 | EPI_ISL_434692 |
| T95I | BA.1, BA.3, and BA.1.1 | Many Delta sublineages, Iota, Mu [37, 38] | Asia / Lebanon | 2020-01-10 | EPI_ISL_8311730 |
| G142D | BA.1, BA.2, BA.3, and BA.1.1 | Delta [36] | Asia / Japan / Kanto | 2020-02-25 | EPI_ISL_479852 |
| V143del | BA.1, BA.3, and BA.1.1 | Alpha, Eta [38] | Asia / Japan / Kanto | 2020-02-25 | EPI_ISL_479852 |
| Y144del | BA.1, BA.3, and BA.1.1 | Alpha, Eta, Mu (Y144S) [37] | Europe / Switzerland / Aargau | 2020-01-05 | EPI_ISL_896091.3 |
| Y145del | BA.1, BA.3, and BA.1.1 | Alpha, Eta, Mu (Y145N) [37, 38] | North America / USA / Virginia | 2020-03-30 | EPI_ISL_429978 |
| N211I | BA.1, BA.3, and BA.1.1 | — | North America / USA / Texas / Houston | 2020-07-14 | EPI_ISL_5343378 |
| L212del | BA.1, BA.3, and BA.1.1 | — | Europe / Spain / Navarra | 2020-05-12 | EPI_ISL_1382928 |
| V213G | BA.2 | — | Asia / Iran | 2020-03-29 | EPI_ISL_672575 |
| ins214EPE | BA.1 and BA.1.1 | — | Africa / Zambia / Lusaka / Lusaka | 2021-02-12 | EPI_ISL_7988145 |
| **Spike protein (RBD)** |  |  |  |  |  |
| G339D | BA.1, BA.2, BA.3, and BA.1.1 | — | Europe / Norway | 2020-03-05 | EPI_ISL_420138 |
| R346K | BA.1.1 | Mu [36] | North America / Panama / Cocle | 2020-03-18 | EPI_ISL_496747 |
| S371L | BA.1 and BA.1.1 | — | North America / USA / Louisiana / Lincoln Parish | 2020-12-27 | EPI_ISL_8508546 |
| S371F | BA.2 and BA.3 | — | Europe / Italy / Campania | 2020-05-17 | EPI_ISL_6832115 |
| S373P | BA.1, BA.2, BA.3, and BA.1.1 | — | North America / USA / Texas / Houston | 2020-06-04 | EPI_ISL_545554 |
| S375F | BA.1, BA.2, BA.3, and BA.1.1 | — | Africa / Egypt / Cairo | 2020-06-08 | EPI_ISL_1097025 |
| T376A | BA.2 | — | Asia / Japan / Saitama | 2021-06-09 | EPI_ISL_3198900 |
| D405N | BA.2 and BA.3 | — | Europe / Portugal | 2021-02-09 | EPI_ISL_1116795 |
| R408S | BA.2 | — | Europe / Netherlands / Zuid-Holland | 2020-03-27 | EPI_ISL_422951 |
| K417N | BA.1, BA.2, BA.3, and BA.1.1 | Beta, Gamma (K417T), two sub lineages of the Delta variant (AY.1 and AY.2) [38] | Asia / Qatar / Doha | 2020-03-27 | EPI_ISL_2274264 |
| **RBD(RBM)** |  |  |  |  |  |
| N440K | BA.1, BA.2, BA.3, and BA.1.1 | — | North America / Canada / Quebec | 2020-03-10 | EPI_ISL_3458197 |
| G446S | BA.1, BA.3, and BA.1.1 | — | Europe / United Kingdom / England | 2020-04-04 | EPI_ISL_440772 |
| S477N | BA.1, BA.2, BA.3, and BA.1.1 | — | Asia / Lebanon | 2020-01-10 | EPI_ISL_8317205 |
| T478K | BA.1, BA.2, BA.3, and BA.1.1 | Delta [38] | North America / USA / New York / Westchester County | 2020-03-12 | EPI_ISL_6661895 |
| E484A | BA.1, BA.2, BA.3, and BA.1.1 | Beta (E484K), Gamma (E484K), Eta (E484K), Iota (E484K), Mu (E484K), Kappa (E484Q) [36, 37] | Europe / Spain / La Rioja | 2020-03-03 | EPI_ISL_578200 |
| Q493R | BA.1, BA.2, BA.3, and BA.1.1 | — | Europe / United Kingdom / England | 2020-04-22 | EPI_ISL_470150 |
| G496S | BA.1 and BA.1.1 | — | Africa / Egypt | 2020-06-06 | EPI_ISL_8194875 |
| Q498R | BA.1, BA.2, BA.3, and BA.1.1 | — | North America / USA / Louisiana / Lincoln Parish | 2020-12-27 | EPI_ISL_8508546 |
| N501Y | BA.1, BA.2, BA.3, and BA.1.1 | Alpha, Beta, Gamma, Mu [37] | Europe / Spain / Castilla y Leon | 2020-02-07 | EPI_ISL_1603195 |
| Y505H | BA.1, BA.2, BA.3, and BA.1.1 | — | Europe / United Kingdom / England | 2020-04-17 | EPI_ISL_465212 |
| **Spike protein (SD1/SD2)** |  |  |  |  |  |
| T547K | BA.1 and BA.1.1 | — | North America / USA / Florida | 2020-03-20 | EPI_ISL_1379457 |
| D614G | BA.1, BA.2, BA.3, and BA.1.1 | Alpha, Beta, Gamma, Delta, Kappa, Eta, Iota, Lambda, Mu [37] | Africa / Niger / Niamey | 2020-01-01 | EPI_ISL_3804266 |
| H655Y | BA.1, BA.2, BA.3, and BA.1.1 | Gamma [37] | Europe / France / Île-de-France / Paris | 2020-01-23 | EPI_ISL_1301441 |
| **Spike protein (PBCS)** |  |  |  |  |  |
| N679K | BA.1, BA.2, BA.3, and BA.1.1 | — | North America / USA / New York / Brooklyn | 2020-03-18 | EPI_ISL_421580 |
| N764K | BA.1, BA.2, BA.3, and BA.1.1 | Alpha, Theta, Mu, Delta (P681R), Kappa (P681R), A.23.1 (P681R) [37, 38] | Africa / Niger / Niamey | 2020-01-01 | EPI_ISL_3804266 |
| **Spike protein (Others CTD)** |  |  |  |  |  |
| N764K | BA.1, BA.2, BA.3, and BA.1.1 | — | Africa / Egypt / Cairo | 2020-06-08 | EPI_ISL_1097025 |
| D796Y | BA.1, BA.2, BA.3, and BA.1.1 | — | Europe / United Kingdom / Scotland | 2020-03-29 | EPI_ISL_433364 |
| N856K | BA.1 and BA.1.1 | — | Asia / China / Hubei / Wuhan | 2020-01-25 | EPI_ISL_493156 |
| Q954H | BA.1, BA.2, BA.3, and BA.1.1 | — | Africa / South Africa / North West | 2020-06-29 | EPI_ISL_8182884 |
| N969K | BA.1, BA.2, BA.3, and BA.1.1 | — | North America / USA / Maryland | 2020-05-31 | EPI_ISL_3098538 |
| L981F | BA.1 and BA.1.1 | — | North America / USA / Wisconsin / Dane County | 2020-04-17 | EPI_ISL_450716 |
| **ORF3a (NS3a)** |  |  |  |  |  |
| T223I | BA.2 and BA.3 | — | Asia / India / Himachal Pradesh | 2020-02-24 | EPI_ISL_2099688 |
| **E protein** |  |  |  |  |  |
| T9I | BA.1, BA.2, BA.1.1, and BA.3 | — | North America / USA / Washington | 2020-03-12 | EPI_ISL_476911 |
| **M protein** |  |  |  |  |  |
| D3G | BA.1 and BA.1.1 | — | Asia / China / Guangxi / Nanning | 2020-02-19 | EPI_ISL_6951243 |
| Q19E | BA.1, BA.2, BA.1.1, and BA.3 | — | North America / USA / New York / Brooklyn | 2020-03-27 | EPI_ISL_456032 |
| A63T | BA.1, BA.2, BA.1.1, and BA.3 | — | Europe / United Kingdom / Scotland | 2020-03-23 | EPI_ISL_425915 |
| **ORF6** |  |  |  |  |  |
| D61L | BA.2 | — | N/A | N/A | N/A |
| **N protein** |  |  |  |  |  |
| P13L | BA.1, BA.2, BA.1.1, and BA.3 | Lambda [36] | Oceania / Papua New Guinea | 2020-01-13 | EPI_ISL_1307662 |
| E31del | BA.1, BA.2, BA.1.1, and BA.3 | — | Africa / Egypt / Cairo | 2020-06-06 | EPI_ISL_1167190 |
| R32del | BA.1, BA.2, BA.1.1, and BA.3 | — | Africa / Egypt / Cairo | 2020-06-06 | EPI_ISL_1167190 |
| S33del | BA.1, BA.2, BA.1.1, and BA.3 | — | Africa / Egypt / Cairo | 2020-06-06 | EPI_ISL_1167190 |
| R203K | BA.1, BA.2, BA.1.1, and BA.3 | Alpha, Gamma, Lambda [36] | Oceania / Australia / New South Wales / Sydney | 2020-01-08 | EPI_ISL_3568416 |
| G204R | BA.1, BA.2, BA.1.1, and BA.3 | Alpha, Gamma, Lambda [36] | Oceania / Australia / New South Wales / Sydney | 2020-01-08 | EPI_ISL_3568416 |
| S413R | BA.2 and BA.3 | — | Africa / Egypt | 2020-06-24 | EPI_ISL_2380075 |

The different Omicron lineages (BA.1, BA.2 and BA.3) share 33 mutations (ORF1a:T3255I, ORF1a:P3395H, ORF1a:Δ3675/3677, ORF1b:P314L, ORF1b:I1566V, S:G142D, S:G339D, S:S373P, S:S375F, S:K417N, S:N440K, S:S477N, S:T478K, S:E484A, S:Q493R, S:Q498R, S:N501Y, S:Y505H, S:D614G, S:H655Y, S:N679K, S:P681H, S:N764K, S:D796Y, S:Q954H, S:N969K, E:T9I, M:Q19E, M:A63T, N:P13L, N:Δ31/33, N:R203K, N:G204R). On the other hand, 15 mutations are unique to their lineages. Most of them (13 mutations) corresponds to the BA.2 lineage (ORF1a:T842I, ORF1a:L3027F, ORF1a:T3090I, ORF1a:L3201F, ORF1a:F3677L, ORF1b:R1315C, ORF1b:T2163I, S:T19I, S:LPPA24Sdel, S:V213G, S:T376A, S:R408S, ORF6:D61L). BA.3 possesses a unique mutation in NSP6:A88V. And only 1 mutation is distinctive to the sub-lineage BA.1.1 (S:R346K). It is notable that the BA.3 lineage has mutations shared with both BA.1 (and its BA.1.1 sub-lineage) and BA.2. However, it is closer to the BA.1 and BA.1.1 because it possesses S:Δ69/70, S:Δ143/145 and S:NL211I deletions, which are not present in BA.2. Detailed annotation of the positions of Omicron mutations are shown in Supplementary Table 1.

### The occurrence of mutation clusters in the previous variants

To see if any mutations appear together in earlier variants we built a Sankey diagram based on the data in Table 1, as shown in Fig 2. We found that all mutations specific to the BA.2 lineage are unique to the Omicron variant. However, mutations specific to BA.1 variant as well as mutations shared between BA.1 and BA.2 have been found in several variants. The BA.1.1 and BA.3 lineages was excluded as BA.1.1 is identical to BA.1 in all mutations except for one mutation that it shares with the Mu variant, and all BA.3 mutations are shared with other lineages.

**Fig 2.**
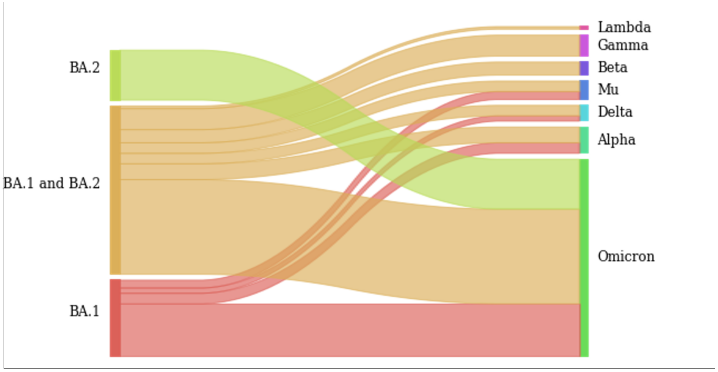
Sankey diagram. This diagram illustrates relationships between different Omicron lineages and variants graphically. The bars on the left show the lineages and the bars on the right represent the variants. The lines between the bars are reflecting the occurrence of mutations. The wider the connection lines are, the more appearance of mutations exists.

### Analysis of the Omicron’s deletions sites for non deletion mutations in other variants

Concerning non-Spike and Spike deletions in Omicron lineages, no other types of mutations were detected in earlier VOCs/VOIs at the same deletion sites, except for PPA25 deletion mutation. In the Spike protein, the P26S substitution mutation has been identified in the Gamma variant, which occurs in the deletion site (25-27del) of the Omicron variant (BA.2 lineage). This substitution also occurs near the NTD supersite at residue with high antibody accessibility scores [39]. This mutation has been linked with reduced binding with monoclonal sera [40].

### Detailed description of each mutation based on published data

#### Mutations in Spike protein

The Spike protein is heavily mutated in the Omicron lineages BA.1, BA.2, BA.3, and BA.1.1 compared to other VOCs and VOIs (Fig 3). There are 21 mutations common in all of these four lineages. Comparing BA.1 and its sub-lineage BA.1.1, Spike protein revealed one additional mutation in BA.1.1 (R346K). There are 37 mutations in the Spike protein of BA.1, 31 mutations in the Spike protein of BA.2, 33 mutations in the Spike protein of BA.3, and 38 mutations in the Spike protein of BA.1.1. Comparing BA.1 and BA.2 lineages, Spike protein revealed 16 specific mutations in BA.1 lineage and 10 specific mutations in BA.2. BA.3 shares 10 mutations from BA.1-specific mutations and 2 mutations from BA.2-specific mutations to form its spike protein.

**Fig 3.**
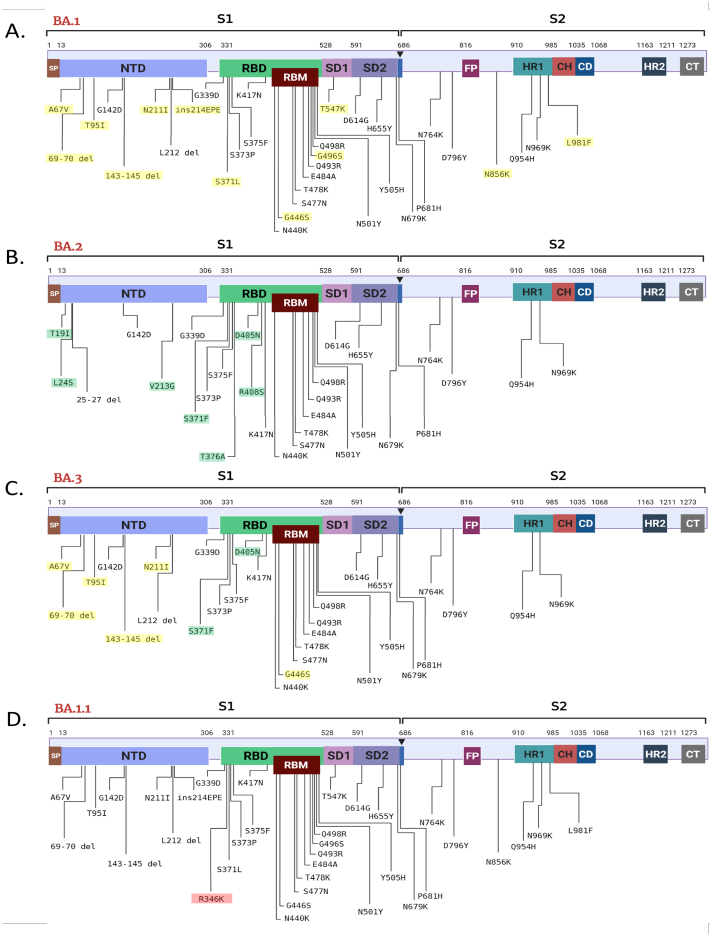
Schematic representation of the Spike protein mutations identified in the genome of the Omicron BA.1, BA.2, and BA.3 lineages, and BA.1.1 sub-lineage. Unique mutations are highlighted in yellow for BA.1 (A), in green for BA.2 (B) and in red for BA.1.1 (D). For BA.3 mutations shared with BA.1 are highlighted in yellow and mutations shared with BA.2 are highlighted in green (C). The Spike protein is composed of two subunits, S1 and S2. Arrow indicates the S1/S2 protease cleavage site. Different domains are shown by different colors. SP: signal peptide; NTD: N-terminal domain; RBD: receptor-binding domain; RBM: receptor-binding motif; SD1: subdomain 1; SD2: subdomain 2; FP: fusion peptide; HR1: heptad repeat 1; CH: central helix; CD: connector domain; HR2: heptad repeat 2; CT, cytoplasmic tail. Created with BioRender [23].

##### 0.1 Mutations in RBD

Mutations in RBD may have a high impact on coronavirus fitness, as this region interacts with hACE2 receptors on human cells. Moreover a combination of few mutations may lead to cumulative effect. For instance, Nelson G et al.showed increased affinity for the ACE2 protein in presence of a combination of K417N, E484K and N501Y mutations, as they significantly affect RBD conformation [41]. But in the case of the Omicron, there is E484A substitution for which it showed the ability to escape neutralising antibodies [42]. Same results were observed for a combination of S477N, E484K and N501Y mutations [43]. Below each mutation is discussed in further detail.

###### 0.1.1 G339D [BA.1, BA.1.1, BA.2, BA.3]

This mutation might decrease binding affinity to ACE2 [44].

###### 0.1.2 R346K [BA.1.1]

Evaluation of the free energy perturbation (FEP) has shown that the effect of this mutation on antibody binding may not be significant, and it only affects class 2 mAbs. However, in the presence of other mutations (e.g. K417N), the neutralising activity of the antibodies may be reduced more significantly [45]. R346K is also found in the Mu variant of interest that emerged in early 2021.

###### 0.1.3 S371F [BA.2, BA.3]

Previous mutation variants found in this position include S371P, S371F, S371T and S371A [46]. The S371 residue is located in one of the cryptic binding epitopes that are aligned with conserved hinge positions. In addition, the S371-F541 region shows a lack of glycosylated sites. Identification of such regions is necessary to search for potential epitopes for neutralising antibodies [47]. On the other hand, changes in the amino acid residue at the S371 position result in reduced RBD recognition by antibodies [48]. For example, a single S371L mutation (which occurs in BA.1) at this position results in avoidance of some neutralising antibodies (by changing the conformation). Possibly, the S371F mutation leads to a similar effect [49, 50].

###### 0.1.4 S371L [BA.1, BA.1.1]

This mutation decreases stability of the RBD and may reduce the ability of antibodies to recognise the Spike protein epitope [49, 51].

###### 0.1.5 S373P [BA.1, BA.1.1, BA.2, BA.3]

This mutation may help to escape neutralising antibodies [52].

###### 0.1.6 S375F [BA.1, BA.1.1, BA.2, BA.3]

This mutation probably first occurred in the Omicron. As it has effects on replacement of serin for phenylalanine, it may slightly change RBD conformation.

###### 0.1.7 T376A [BA.2]

The T376 residue is located in the 2-strand, which may be an epitope for neutralising antibodies [53]. This mutation may be involved in evading recognition by neutralising antibodies, similar to other mutations in RBD [49].

###### 0.1.8 D405N [BA.2, BA.3]

The D504N/Y mutation results in the escape of one of the monoclonal antibody groups that bind to the RBD base [54]. An important feature of this mutation is that it is located close to RBM, specifically the G504 residue. Hence can affect the RBM conformation [55]. In addition, residue D405 is involved in the Spike protein transition to the open form [56].

###### 0.1.9 R408S [BA.2]

The R408K mutation in this position has a similar effect to the D504N mutation in reducing epitope recognition in RBD by monoclonal antibodies [54].

R408 as well as D405 are involved in the RBD-opening transition [56]. Thus, the study of these residues is important to elucidate their role for viral entry into the cell and how mutations affect this ability.

###### 0.1.10 K417N [BA.1, BA.1.1, BA.2, BA.3]

This mutation helps the virus to escape monoclonal antibodies [57]. This mutation alone showed a slightly lower or similar affinity for ACE 2 as wild type [58].

###### 0.1.11 N440K [BA.1, BA.1.1, BA.2, BA.3]

Based on binding-free energy change, this mutation potentially might increase binding affinity to ACE2 receptors [44].

###### 0.1.12 G446S [BA.1, BA.1.1, BA.3]

While reducing RBD stability it is increasing ACE2 binding affinity [59]. Might help to escape neutralising antibodies [60].

###### 0.1.13 S477N [BA.1, BA.1.1, BA.2, BA.3]

This mutation shows resistance to neutralising monoclonal antibodies and increased ACE2 binding [8, 61]. Hence this mutation provides stronger transmissibility. At the same time, this mutation showed potency to lower binding affinity [44].

###### 0.1.14 T478K [BA.1, BA.1.1, BA.2, BA.3]

This mutation arose multiple times independently. Since it occurs with high frequency, it may positively affect ACE2 binding affinity [62].

###### 0.1.15 E484A [BA.1, BA.1.1, BA.2, BA.3]

This mutation abolished binding of some neutralising antibodies [42, 63].

###### 0.1.16 Q493R [BA.1, BA.1.1, BA.2, BA.3]

This substitution reduces protein’s susceptibility to some monoclonal antibodies in neutralisation assay [64].

###### 0.1.17 G496S [BA.1, BA.1.1]

G496 is one of the key residues that interact with ACE2. For substitutions G496W and G496Y showed the largest binding energy change among all RBD mutations [65]. G496S substitution has the same effect of enhancing binding capacity [66].

###### 0.1.18 Q498R [BA.1, BA.1.1, BA.2, BA.3]

In combination with N501Y increases affinity to ACE2 [43].

###### 0.1.19 N501Y [BA.1, BA.1.1, BA.2, BA.3]

This mutation appeared multiple times independently. It may affect affinity resulting in increasing ACE2 binding [41]. Key mutation in combination K417N + E484K + N501Y [57].

###### 0.1.20 Y505H [BA.1, BA.1.1, BA.2, BA.3]

Chen et al. showed that Y505 residue is a potential hotspot and has a most likely substitution Y505H [61]. Nevertheless this mutation may lead to reducing ACE binding affinity [59].

##### 0.2 Mutations in other Spike protein regions

###### 0.2.1 T19I [BA.2]

Variants of mutations in this position for the first half year of the pandemics were T19P, T19I, T19S [67]. But the Delta variant already contained a T19R substitution, hence it is currently the most common [68]. The T19 site is located in the NTD in a region with a very high mutation density. As the NTD is located close to the RBD, it can therefore be an epitope for antibodies [69, 70]. Thus, it can be assumed that this high variability of this site is a mechanism of antibody recognition avoidance.

###### 0.2.2 LPPA24S [BA.2]

This mutation results in a nucleotide deletion at positions 21633-21641 (TACCCCCTG), which causes a replacement LPPA24S. It probably has the same effect as the previous mutation, consisting in the evasion of recognition by neutralising antibodies.

###### 0.2.3 A67V [BA.1, BA.1.1, BA.3]

Substitution of Ala for Val results in forming of novel hydrophobic bonds with I100, F79 and A263 residues. Therefore it slightly changes conformation of 3-4 loops [69].

###### 0.2.4 del69-70 [BA.1, BA.1.1, BA.3]

This mutation might allosterically change S1 conformation and be supportive for mutations in RBD [71, 72]. These deletions are associated with increased virus replication and also involved in the evasion of neutralizing antibodies [30, 73].

###### 0.2.5 T95I [BA.1, BA.1.1, BA.3]

One of the most prevalent mutations in the Delta variant [74]. Protein modeling has shown that by altering the NTD 3D structure may enhance G142 mutation and hence increase viral load [75].

###### 0.2.6 G142D [BA.1, BA.1.1, BA.2, BA.3] + Δ143/145 [BA.1, BA.1.1, BA.3]

Similar deletion (del141-144) was found in isolated culture on 70th day obtained from immunocompromised patient with cancer [76]. Because of the location of this deletion in the recurrent deletion region 2 (RDR2), it can affect the ability to escape neutralising antibodies [77].

###### 0.2.7 Y145D

Spike protein with this mutation has shown ability to escape some neutralising antibodies [78].

###### 0.2.8 NL211I del [BA.1, BA.1.1, BA.3]

This mutation locates in the recurrent deletion region 3 (RDR3) and may affect ability to escape some neutralising antibodies [77].

###### 0.2.9 V213G [BA.2]

This mutation locates in the NTD of the Spike protein. No significant effects have been shown for it.

###### 0.2.10 ins214EPE [BA.1, BA.1.1]

86.3% of the Omicron variant samples contain this insertion (nucleotides GAGCCA-GAA) [79]. Like other alternative insertions in this site, it may affect SARS-CoV-2 infectivity as located next to RDR3 [80].

###### 0.2.11 T547K [BA.1, BA.1.1]

Cases of this mutation were already registered [52], but it was first fixed in the Omicron.

###### 0.2.12 D614G [BA.1, BA.1.1, BA.2, BA.3]

In March of 2020, this mutation first occurred, and very fast became prevalent in most samples. Facilitate proteolysis in furin cleavage site and hence may increase transmissibility [81].

###### 0.2.13 H655Y [BA.1, BA.1.1, BA.2, BA.3]

Braun, K. et al. suggested that this mutation might be under positive selection in domestic cats. Associated with furin cleavage site [82].

###### 0.2.14 N679K and P681H [BA.1, BA.1.1, BA.2, BA.3]

Both these mutations are located in the region of S1/S2 junction. Changing from neutral charged amino acids to the positive charged residues results in enhancing of coupling furin cleavage site with proteolytic enzyme hence that leads to increasing cleavability of spike S1/S2 site [83]. But at the same time, for a single N679K substitution, it is shown that it alone is not enough to increase transmissibility [84].

###### 0.2.15 N764K [BA.1, BA.1.1, BA.2, BA.3]

This mutation first fixed in the Omicron. It might decrease protein stability and reduce its function [51].

###### 0.2.16 D796Y [BA.1, BA.1.1, BA.2, BA.3]

Appeared a few times independently [30, 85], including a case of intra-host evolution in a immunocompromised (HIV) patient [27].

###### 0.2.17 N856K [BA.1, BA.1.1]

Located between fusion peptide (FP) and heptad repeat 1 (HR1) regions of Spike protein [86]. The mutation consists of a substitution with a positively charged amino acid, which can locally affect the conformation of this region and increase the affinity to certain proteins.

###### 0.2.18 Q954H, N969K [BA.1, BA.1.1, BA.2, BA.3] and L981F [BA.1, BA.1.1]

These mutations are located within the heptad repeat 1 (HR1) region that acts in the process of fusion with the host cell. Thus these mutations may affect transmissibility and infectivity [86, 87].

## 1 Non-Spike mutations

Non-Spike regions are remarkably mutated in the Omicron lineages BA.1, BA.2, BA.3, and BA.1.1 compared to other VOCs and VOIs (Fig 4). There are 16 non-Spike mutations common in all of these four lineages. Comparing BA.1 and its sub-lineage BA.1.1, revealed that there is no additional mutations in the non-Spike proteins of BA.1.1. Comparing BA.1 and BA.2 lineages, there are 6 specific mutations in BA.1 lineage and 11 specific mutations in BA.2. BA.3 shares 5 mutations from BA.2-specific mutations and has one specific mutation (NSP6:A88V).

**Fig 4.**
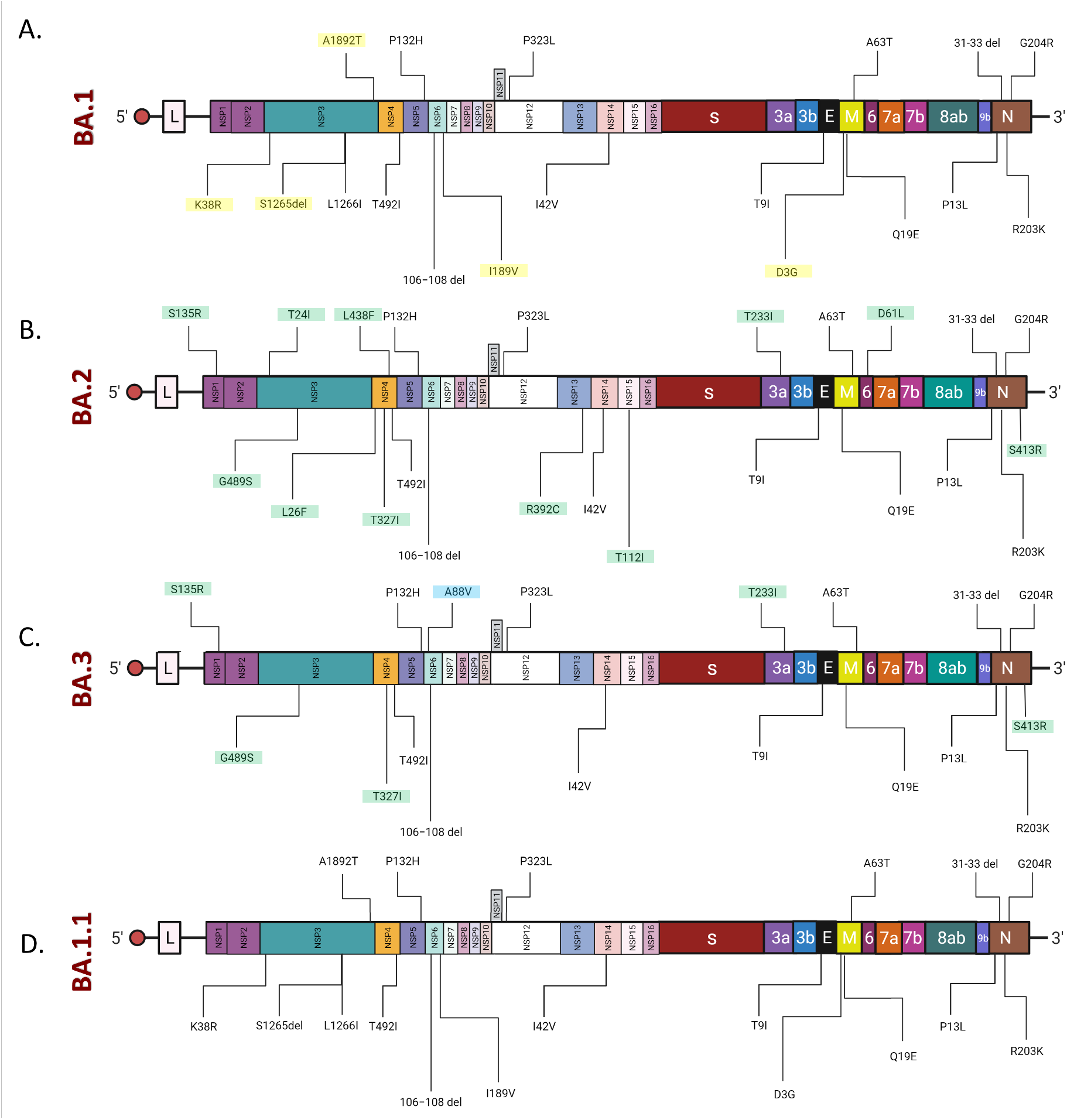
Schematic representation of the non-Spike mutations identified in the genomes of the Omicron BA.1, BA.2 and BA.3 lineages, and BA.1.1 sub-lineage. Unique mutations are highlighted in yellow for BA.1 (A), in green for BA.2 (B) and in blue for BA.3 (C). Mutations shared between BA.2 and BA.3 are highlighted in green (C). There are 22 mutations outside the Spike protein of BA.1, 27 mutations outside the Spike protein of BA.2, 22 mutations outside the Spike protein of BA.3, and 22 mutations outside the Spike protein of BA.1.1. Created with BioRender [23].

### 1.1 Mutations in E protein

#### 1.1.1 T9I [BA.1, BA.1.1, BA.2, BA.3]

Located in the transmembrane domain, this mutation can affect the configuration of E protein and provides a stronger anchoring to the viral membrane [88, 89].

### 1.2 Mutations in M protein

#### 1.2.1 D3G [BA.1, BA.1.1]

One of the most frequent mutations in M protein [90]. Locates in the exposed NTD region, mutation in which may affect the interactions with host cells [91].

#### 1.2.2 Q19E and A63T [BA.1, BA.1.1, BA.2, BA.3]

There is no data on effects of these mutations yet.

### 1.3 Mutations in N protein

#### 1.3.1 P13L [BA.1, BA.1.1, BA.2, BA.3]

Oulas, A. et al. suggested that this mutation is associated with lower levels of transmissibility and death rates [92].

#### 1.3.2 Δ31-33 [BA.1, BA.1.1, BA.2, BA.3]

Deletion consists in the substitution of “GERS” for “G”. Located near the N-terminal domain, thus can affect the assembly with M protein [93].

#### 1.3.3 R203K and G204R [BA.1, BA.1.1, BA.2, BA.3]

R203K and G204R mutations in nucleocapsid have been shown to be linked to increased subgenomic RNA expression [94] and increased viral loads [95]. In addition, these mutations are found in patients with inferior outcomes [96].

#### 1.3.4 S413R [BA.2, BA.3]

N protein is highly phosphorylated and enriched with basic residues. Basic residues properties help to form a sustainable complex with viral RNA, while the role of phosphorylation is not defined. However, for phosphorylation null mutants viral particles have been shown to be assembled with comparable or slightly lower efficiency than the wild type [97]. Therefore, the loss of phosphorylation at some sites might negatively affect the process of packaging RNA into the nucleocapsid envelope. Residue S413 is one of the phosphorylation sites. From one side mutation S413R changes polar uncharged Serine for basic charged Arginine, on another side it removes phosphorylation from Serine. Hence it is unclear whether the S413R mutation has a positive effect on viral particle assembly (change to the major residue) or a negative effect (it eliminates phosphorylation).

### 1.4 Mutations in NSP1 (leader protein)

#### 1.4.1 S135R [BA.2, BA.3]

Hydrogen-bond interaction analysis showed that S135 residue in NSP1 forms hydrogen bonds with C16 and C20 nucleotides of SARS-CoV-2 5’-UTR stem loop [98]. Substitution of Serine for Arginine may increase RNA compatibility and as result enhance recognition.

### 1.5 Mutations in NSP3

#### 1.5.1 T24I (ORF1ab: T842I) [BA.2]

This mutation first fixed in B.1.1.529. No significant effects have been reported for it, yet. It locates in the ubiquitin-like domain 1 [99].

#### 1.5.2 K38R (ORF1ab: K856R) [BA.1, BA.1.1]

Locates in the ubiquitin-like domain 1 [99].

#### 1.5.3 G489S (ORF1ab: G1307S) [BA.2, BA.3]

No effects have been reported for this mutation yet.

#### 1.5.4 1265 + L1266I (ORF1ab: SL2083I) [BA.1, BA.1.1]

No effects have been reported for these mutations yet.

#### 1.5.5 A1892T (ORF1ab: A2710T) [BA.1, BA.1.1]

No significant effects have been reported for this mutation yet.

### 1.6 Mutations in NSP4

#### 1.6.1 L264F (ORF1ab: L3027F) [BA.2]

This mutation may have appeared several times independently in different lines [100, 101].

#### 1.6.2 T327I (ORF1ab: T3090I) [BA.2]

This mutation was detected in some samples early in the pandemic [102, 103], but has not become widespread since then.

#### 1.6.3 L438F (ORF1ab: L3201F) [BA.2]

L438F mutation was detected several times before, but has not become widespread [104].

#### 1.6.4 T492I (ORF1ab: T3255I) [BA.1, BA.1.1, BA.2, BA.3]

One of the most frequently co-occurring with S:T478K non-spike mutation [62].

### 1.7 Mutations in NSP5

#### 1.7.1 P132H (ORF1ab: P3395H) [BA.1, BA.1.1, BA.2, BA.3]

No significant effects have been noted for this mutation. The crystal structure shows that it is located away from the active centre and both alternative binding regions [105].

### 1.8 Mutations in NSP6

#### 1.8.1 A88V (ORF1ab: A3657V) [BA.3]

No effects have been reported for this mutation yet.

#### 1.8.2 L105F (ORF1ab:L3674F) [BA.1, BA.1.1, BA.2] + Δ105-107 (ORF1ab: SGF3675del) [BA.1, BA.1.1, BA.2, BA.3]

This deletion results in the replacement of amino acids ‘SLSG’ by ‘S’ (Serine). This Serine is directly followed by Phenylalanine, for which it has been shown to increase the affinity of NSP6 to the endoplasmic reticulum membrane. It is suggested that this binding helps to avoid the delivery of viral particles to the lysosomes. Hence, this deletion may be part of the mechanism of innate immunity evasion [106, 107].

#### 1.8.3 F108L (ORF1ab: F3677L) [BA.2]

Is more likely to occur in P.1 (18.2%) [108], but globally the deletion (NSP6:S106/G107/F108del) is more common in this position. According to GISAID (as on 1 September 2021), this deletion is found in 97.21% of the sequences [109].

#### 1.8.4 I189V (Orf1ab: I3758V) [BA.1, BA.1.1]

Sun et al. suggested that this mutation may affect viral shedding and hence transmissibility, thereby altering viral load [110]. Although this assumption requires experimental verification.

### 1.9 Mutations in NSP12

#### 1.9.1 P323L (ORF1ab: P4715L) [BA.1, BA.1.1, BA.2, BA.3]

At least in Turkey this mutation has been found in a large number of samples [111] and presumably in combination with D614G mutation in the spike protein may have a correlation with severity of the disease [112].

### 1.10 Mutations in NSP13

#### 1.10.1 R392C (ORF1ab: R5716C) [BA.2]

Sengupta et al. analysed the effects of different mutations on the properties and functions of coronavirus proteins. PROVEAN (Protein Variation Effect Analyzer) score showed that R392C mutation have deleterious effects on NSP13 protein and Gibbs free energy (ΔΔG) measured -1.29 [113]. In addition, this mutation results in the substitution of a positively charged amino acid for a polar uncharged residue. Thus, this mutation may lead to a slight change in protein folding. The [114] study also suggests the ability of this mutation to alter the secondary structure of the protein.

In humans, the R392C mutation is less frequent (*<*0.5% of human isolates) and is more common in the mink population (10.5% of mink isolates) [115].

### 1.11 Mutations in NSP14

#### 1.11.1 I42V (ORF1ab: I5967V) [BA.1, BA.1.1, BA.2, BA.3]

The I42V mutation is not expected to have a significant effect on the protein structure as it consists of the substitution of one hydrophobic amino acid (Isoleucine) for another hydrophobic amino acid (Valine).

### 1.12 Mutations in NSP15

#### 1.12.1 T112I (ORF1ab: T6564I) [BA.2]

No effects have been reported for this mutation yet.

### 1.13 Mutations in ORF3a

#### 1.13.1 T223I [BA.2, BA.3]

Located in the cytosolic domain near the hydrophobic core. Replacing the polar uncharged amino acid (Threonine) with a hydrophobic amino acid (Isoleucine) can slightly change the protein conformation [116].

### 1.14 Mutations in ORF6

#### 1.14.1 D61L [BA.2]

The D61L mutation results in the substitution of a negatively charged residue (Aspartic acid) with a hydrophobic residue (Leucine). This substitution can lead to the formation of a stronger ORF6:IRF3 complex. This in turn leads to an increase in the antagonistic properties of ORF6 against IFN [88].

## 2 Spike mutations aren’t the only ones that matter

### Mutations in NSP1 are important

NSP1 (or leader protein) interacts with 40S subunit of the ribosome, inhibits host gene expression, and also evades the immune system. Furthermore, NSP1 degrades host mRNA and facilitates viral gene expression [117, 118]. Leader protein contains a globular region that recognize the stem loop of the 5’-UTR region of SARS-CoV-2 RNAs hence viral genes continue to translate [98].

This protein has been proposed as a target for vaccine development and also for drug design. The majority of mutations occuring in NSP1 exhibited a destabilizing effect and increased flexibility [119].

### Mutations in NSP3 are important

NSP3 has been proposed to work with nsp4 and nsp6 to induce double-membrane vesicles (DMVs), which serve as an important component of the replication/transcription complex (RTCs) [120]. Moreover NSP3 interacts with the N-terminal domain of the nucleocapsid phosphoprotein (N), which leads to binding of the nucleocapsid and RTCs [121]. NSP3 also antagonists type I interferon mediated immune response, and blocks NF-kappa-signal transduction [122, 123]. Mutations in NSP3 had been linked with positive selection of viruses leading to evolution in beta-coronaviruses [124].

### Mutations in NSP4 are important

NSP4 participates in assembly of cytoplasmic DMVs and helps in viral replication [120, 125].

### Mutations in NSP5 are important

NSP5 or main protease (Mpro), also known as 3C-like protease (3CLpro) is essential for viral life cycle as it cleaves polyproteins pp1ab and pp1a into distinct functional proteins [126]. Therefore, this protein is a putative target for anticoronavirus therapy. NSP5 may be involved in evading the innate immune response by inhibiting mitochondrial antiviral signalling (MAVS) protein and IFN induction [127]. Mutations in this protein can affect its proteolytic activity.

### Mutations in NSP6 are important

Along with NSP3 and NSP4 is involved in the assembly of DMVs, which contain replication/transcription complexes. Furthermore, NSP6 has also been shown to have affinity to endoplasmic reticulum, which is provided by phenylalanine residues in the sequence of this protein [106].

Deletion mutation in NSP6 from 105-107 could aid in innate immune evasion, possibly by compromising cells’ ability to degrade viral components [106].

### Mutations in NSP12 (RNA-dependent RNA polymerase) are important

Variants of RNA-dependent RNA polymerase (RdRp) emerged early during the COVID-19 outbreak in China, North America, Europe, and Asian countries and hence RdRp was considered as a mutation hotspot [128, 129]. The RdRp:P323L mutation was found to be associated with increasing point mutations in viral isolates in Europe during the early phase of COVID-19 outbreak [130]. Thus, it is possible that mutations in RdRp might alter the interaction of RdRp with the cofactors which could yield less effective proofreading activity leading to the emergence of multiple SARS-CoV-2 variants [128]. In silico analysis predicted the docking site of antiviral drugs within a hydrophobic cleft located near the RdRp:P323L mutation site. This mutation was predicted to diminish the affinity of RdRp for existing antiviral drugs [128].

### Mutations in NSP13 are important

NSP13 has several essential functions, including: (1) helicase activity in the 5’-*>*3’ direction, (2) the ability to unwind RNA/DNA duplex, (3) 5’m-RNA capping activity. Mutations in the active sites of this protein can affect the metabolism of the virus [126, 131].

### Mutations in NSP14 are important

NSP14 acts as a 3’–5’ exoribonuclease (N-terminal domain) for RNA replication proof-reading. In addition, this protein has a second function acting as an N7-methyltransferase (C-terminal domain). Consequently, mutations in this protein may affect the proofreading of newly synthesised viral mRNAs or their stability [132]. Potentially can inhibit interferon signalling [133].

### Mutations in NSP15 are important

NSP15 is an endoribonuclease that cleaves the 3’-end of uridylates [134]. This functionality allows coronaviruses to avoid an innate immune response by cleaving the 5’-polyuridines of the viral RNA, and hence preventing the activation of dsRNA sensors [126, 135].

NSP15 is an essential protein for the viral life cycle, as it has been shown that mutations in this protein can lead to a rapid antiviral response in macrophages, thereby suppressing infection in a short period of time [135].

### Mutations in ORF3a are important

Forms the ion channel. Initiates the lysis process of the host cells to allow new viral particles shedding. Also involved in the autophagy inhibition and disruption of lysosomes. Therefore, it is an essential protein for the viral cycle [116, 131].

Because ORF3a is located on the surface of the membrane, it is able to induce a cellular and humoral immune response in infected individuals [136].

### Mutations in M protein are important

Most conserved protein and most abundant structural protein. Have ability to interact with N, S proteins and viral RNA [90]. Mutations occurring in the M protein could influence the host cell interaction [137].

### Mutations in E protein are important

The mutations found in the E protein may change the structural conformation of this protein and subsequently alter the associated functions such as viral assembly, replication, propagation, and pathogenesis as also previously observed in SARS-CoV [138, 139].

### Mutations in ORF6 are important

It has been shown that ORF6 can act as a suppressor of interferon signalling by inhibiting primary interferon production. ORF6 can therefore interfere in the induction of innate immune response. Molecular docking and dynamics simulation analysis showed that the C-terminal region of the ORF6 protein is able to interact with the transcription factor IRF3 via hydrophobic bonds. Thus, ORF6 acts as an IFN antagonist [88].

### Mutations in N protein are important

Nucleocapsid phosphoprotein interacts with M protein during viral assembling. Has N-terminal and C-terminal domains that can bind RNA. All SARS-CoV-2 variants of interest or concern defined by the WHO contain at least one mutation with *>*50% penetrance within seven amino acids (N:199-205) in the nucleocapsid (N) protein, indicating significance of this region [36, 140].

### 3D Spike Model for Omicron lineages

The 3D structure of reference SARS-CoV-2 spike glycoprotein (closed state) is available in Protein Data Bank (PDB) with PDB ID: 6VXX. Similarly, the reference spike protein domain bound with ACE2 is obtainable with ID 6M0J. The crystalline cryo-EM structure of the Omicron lineage BA.1 spike protein (ID:7T9J) in complex with human ACE2 (ID:7T9K) had been added. However, the crystalline structure of spike protein for BA.1.1, BA.2, and BA.3 are not available.

Although, Parvez S. et al. build a 3D model of spike RBD of SARS-CoV-2 Omicron variant using SWISS-MODEL homology modelling [141] and wild-type spike protein (PDB ID 6M1) as a template. Their model is of good quality and highly reliable. Their validation analysis showed that overall quality factor of the model was 97.5 91.8% residues were in the most favourable regions with 7.6% residues in highly allowed regions [142]. Kumar et al. have generated the structure of BA.1.1, BA.2, and BA.3 by homology modeling with the SWISS-MODEL server. The template PDB ID:7T9L is used to build the 3D structure, and the sequence identity for BA.1.1, BA.2, and BA.3 are 99.50%, 97.01%, and 98.51%, respectively. To evaluate the structure of homology models Ramachandran plot and Errat plot were used. Ramachandran plot of homology modeled RBD of BA.2 and BA.3 showed 90.2% residues in preferred areas and 9.8% residues in additional permitted regions while Errat score for both is 89.4 [21].

The molecular docking between RBD and ACE2 receptor shows that BA.1, BA.1.1, BA.2, and BA.3 have a greater affinity of binding compared to reference SARS-CoV-2 which might cause enhanced transmission of Omicron lineages. At the interface 19 residues of spike protein and 18 residues of ACE-2 protein interact with each other. On the RBD, among the 12 common mutations, 5 showed a higher binding affinity with hACE2, which was also affirmed by previous research conducted by Rehman et al. [143]. We created a 3D model of the spike RBD region using PyMOL software [144]. We labeled only unique mutations for each lineage that are not shared between lineages (Fig 5).

**Fig 5.**
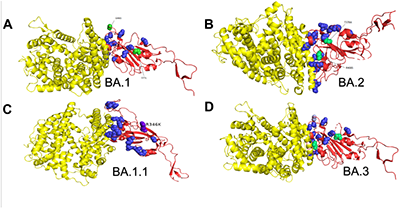
3D modelling of spike RBD of SARS-CoV-2 Omicron lineages. ACE2 in yellow, RBD in red, Mutations shared between all lineages are in blue. Unique mutant residues in the RBD of BA.1 (A) and BA.2 (B) lineages are labeled. (C) The RBD structure of the BA.1.1 sub-lineage with its only one unique mutant residue. (D) The RBD structure of the BA.3 lineage with highlighted mutant residues, Light green: mutation shared between BA.2 and BA.3, Grey: mutation shared between BA.1 and BA.3. Created with PyMOL [144].

## Conclusion

Virological characteristics of newly emerging SARS-CoV-2 variants, such as pathogenicity, transmissibility, resistance to antiviral drugs, and vaccine-induced immunity, are a critical global health concern. Omicron was first reported from South Africa at the end of November 2021. Then the Omicron lineages have rapidly spread worldwide and outcompeted other variants such as Delta. In this study, the mutational spectrum of Omicron lineages (BA.1, BA.1.1, BA.2, and BA.3) was explored in detail. We demonstrated where and when each of these mutations has been previously detected, which mutations were observed in the earlier variants, and how Omicron differs from them. Our study established that the Omicron variant has maintained several mutations that are found in other variants of concern and are thought to make the virus more infectious. We have also found that many of Omicron’s mutations, are extremely rare in other SARS-CoV-2 variants, and represent a considerable jump in SARS-CoV-2 evolution. Accordingly, the mutation rate of the Omicron variant is exceeding the other variants. As well, it has enhanced transmissibility and immune evasion. This progression demonstrates the attempt of the Omicron variant to drive the SARS-CoV-2 for heightened viral fitness.

Based on the Omicron mutation profile in the non-spike regions, we have clarified that it might have collectively enhanced infectivity relative to previous SARS-CoV-2 variants. These mutations have shown to be directly/indirectly associated with the transmission advantages and immune escape. Accordingly, these mutations are worth more studies and investigation. Certainly, more-fit variations can be anticipated to develop over time, and then the occurrence of which has to be monitored thoroughly, as these pose a possible public health threat. The good news is that nothing is infinite, and, eventually, new variants will provide no further advantage in infectivity.

## Supplementary materials

**Supplementary Table 1. Detailed annotation of Omicron mutations**

**Supplementary Table 2. Characteristics of VOCs and VOIs**.

## Data availability

Tables in different formats are available from Github repository: https://github.com/aglabx/omicronic.

## Acknowledgments

We gratefully acknowledge the authors, originating and submitting laboratories where the clinical specimens and/or virus isolates were obtained and published through GISAID on which this research was based. We also thank the BioRender’s authors for their library icons which helps with figure design.

