## Supplementary Table 1. for "Follow-up investigation and detailed mutational characterization of the SARS-CoV-2 Omicron variant lineages (BA.1, BA.2, BA.3 and BA.1.1)"

| Detailed annotation of Omicron mutations | | | | | | | |
|---|---|---|---|---|---|---|---|
| mutation (nuc) | mutation (aa) | BA.1 | BA.1.1 | BA.2 | BA.3* | | |
| T670G | S135R | | | | | NSP1 | |
| C2790T | T842I | | | | | | |
| A2832G | K856R | | | | | | |
| G4184A | G1307S | | | | | NSP3 | |
| 6513_6515del | SL2083I | | | | | | |
| G8393A | A2710T | | | | | | |
| C9344T | L3027F | | | | | | |
| C9534T | T3090I | | | | | | |
| C9866T | L3201F | | | | | NSP4 | ORF1a |
| C10029T | T3255I | | | | | | |
| C10449A | P3395H | | | | | NSP5 | |
| C11235T | A3657V | | | | | | |
| G11287T | L3674F | | | | | | |
| 11288_11296del | SGF3675-3677del | | | | | NSP6 | |
| T11296G | F3677L | | | | | | |
| A11537G | I3758V | | | | | | |
| C14408T | P4715L | | | | | NSP12 | |
| C17410T | R5716C | | | | | NSP13 | ORF1b |
| A18163G | I5967V | | | | | NSP14 | |
| C19955T | T2163I | | | | | NSP15 | |
| C21618T | T19I | | | | | | |
| 21633_21641del | LPPA24S | | | | | | |
| C21762T | A67V | | | | | | |

| | | | | | | | | |
|---|---|---|---|---|---|---|---|---|
| 21765_21770del | HV69-70del | | | | | S gene | NTD | |
| C21846T | T95I | | | | | | | |
| G21987A | G142D | | | | | | | |
| 21987_21995del | VYY143-145del | | | | | | | |
| 22194_22196del | NL211I | | | | | | | |
| T22200G | V213G | | | | | | | |
| 22205GAGCCAGAAins | 214EPEins | | | | | | | |
| G22578A | G339D | | | | | | RBD | |
| G22597A | R346K | | | | | | | |
| C22674T | S371F | | | | | | | |
| T22673C, C22674T | S371L | | | | | | | |
| T22679C | S373P | | | | | | | |
| C22686T | S375F | | | | | | | |
| A22688G | T376A | | | | | | | |
| G22775A | D405N | | | | | | | |
| A22786T | R408S | | | | | | | |
| G22813T | K417N | | | | | | | |
| T22882G | N440K | | | | | | | RBM |
| G22898A | G446S | | | | | | | |
| G22992A | S477N | | | | | | | |
| C22995A | T478K | | | | | | | |
| A23013C | E484A | | | | | | | |
| A23040G | Q493R | | | | | | | |
| G23048A | G496S | | | | | | | |
| A23055G | Q498R | | | | | | | |

| | | | | | | | | |
|---|---|---|---|---|---|---|---|---|
| A23063T | N501Y | | | | | | | |
| T23075C | Y505H | | | | | | | |
| C23202A | T547K | | | | | | **SD1** | |
| A23403G | D614G | | | | | | | |
| C23525T | H655Y | | | | | | **SD2** | |
| T23599G | N679K | | | | | | | |
| C23604A | P681H | | | | | | | **S1/S2** |
| C23854A | N764K | | | | | | | |
| G23948T | D796Y | | | | | | | |
| C24130A | N856K | | | | | | | |
| A24424T | Q954H | | | | | | | |
| T24469A | N969K | | | | | | | |
| C24503T | L981F | | | | | | | |
| C26060T | T223I | | | | | **ORF3a** | | |
| C26270T | T9I | | | | | **E gene** | | |
| A26530G | D3G | | | | | | | |
| C26577G | Q19E | | | | | **M gene** | | |
| G26709A | A63T | | | | | | | |
| G27382C, A27383T, T27384C | D61L | | | | | **ORF6** | | |
| C28311T | P13L | | | | | | | |
| 28362_28370del | ERS31del | | | | | | | |
| G28881A, G28882A | R203K | | | | | **N gene** | | |
| G28883C | G204R | | | | | | | |
| A29510C | S413R | | | | | | | |

* Due to the small number of BA.3 genomes sequenced, there may be inaccuracies in the listed mutations
