## Supplementary Table 2. for "Follow-up investigation and detailed mutational characterization of the SARS-CoV-2 Omicron variant lineages (BA.1, BA.2, BA.3 and BA.1.1)"

The full list of variants and predicted effects

| | Alpha (B.1.1.7) | Beta (B.1.351) | Gamma (P.1) | Delta (B.1.617.2) | Omicron (B.1.1.529) | Lambda (C.37) | Mu (B.1.621) |
|---|---|---|---|---|---|---|---|
| **First detected** | September 2020 | May 2020 | November 2020 | October 2020 | November 2021 | Dec-2020 | January 2021 |
| **Place of detection** | England | South Africa | Brazil | India | South Africa/Netherlands | Peru | Colombia |
| **Number of Mutations** | 23 (17 of which change amino acids) | 21 (8 of which change amino acids) | 17 (11 of which change amino acids) | 12 mutations | ~50 mutations | ~21 mutations including 6 amino acids deletion in the Spike protein | ~21 mutations |
| **Transmissibility** | increased in 29% (95% CI: 24–33) [1] (relative to previously circulating variants at the time and place of emergence) | increased in 25% (95% CI: 20–30) [1] (relative to previously circulating variants at the time and place of emergence) | increased in 38% (95% CI: 29–48) [1] (relative to previously circulating variants at the time and place of emergence) | increased in 97% (95% CI: 76–117) [1] (relative to previously circulating variants at the time and place of emergence) | increased in 105% (95% CI: 96-114) compared with Delta variant [3] | increased compared with the original Wuhan isolate [4] | could have increased tansmissibility [5] |
